# How to overcome the ATRA resistance with the 9aaTAD activation domains in retinoic acid receptors

**DOI:** 10.1101/298646

**Authors:** Martin Piskacek, Marek Havelka, Andrea Knight

## Abstract

In higher metazoa, the nuclear hormone receptors activate transcription trough their specific adaptors, nuclear hormone receptor cofactors NCoA, which are surprisingly absent in lower metazoa. In this study, we demonstrated that the 9aaTAD from NHR-49 receptor activates transcription as a small peptide. We showed, that the 9aaTAD domains are conserved in the human nuclear hormone receptors including HNF4, RARa, VDR and PPARg. The small 9aaTAD peptides derived from these nuclear hormone receptors also effectively activated transcription and that in absence of the NCoA adaptors. We identified adjacent inhibitory domains in the human HNF4 and RARa, which hindered their activation function.

In acute promyelocytic leukaemia (PML-RARa), the receptor mutations often caused all-trans retinoic acid (ATRA) resistance. The fact that almost the entire receptor is needed for ATRA mediated receptor activation, this activation pathway is highly susceptible for loss of function when mutated. Nevertheless in the most of the reported mutants, the activation domains 9aaTAD are still intact. The release of activation 9aaTAD from its dormancy by a new drug might be the sound strategy in combat the ATRA resistance in PML leukaemia.

**Graphical Abstract:** 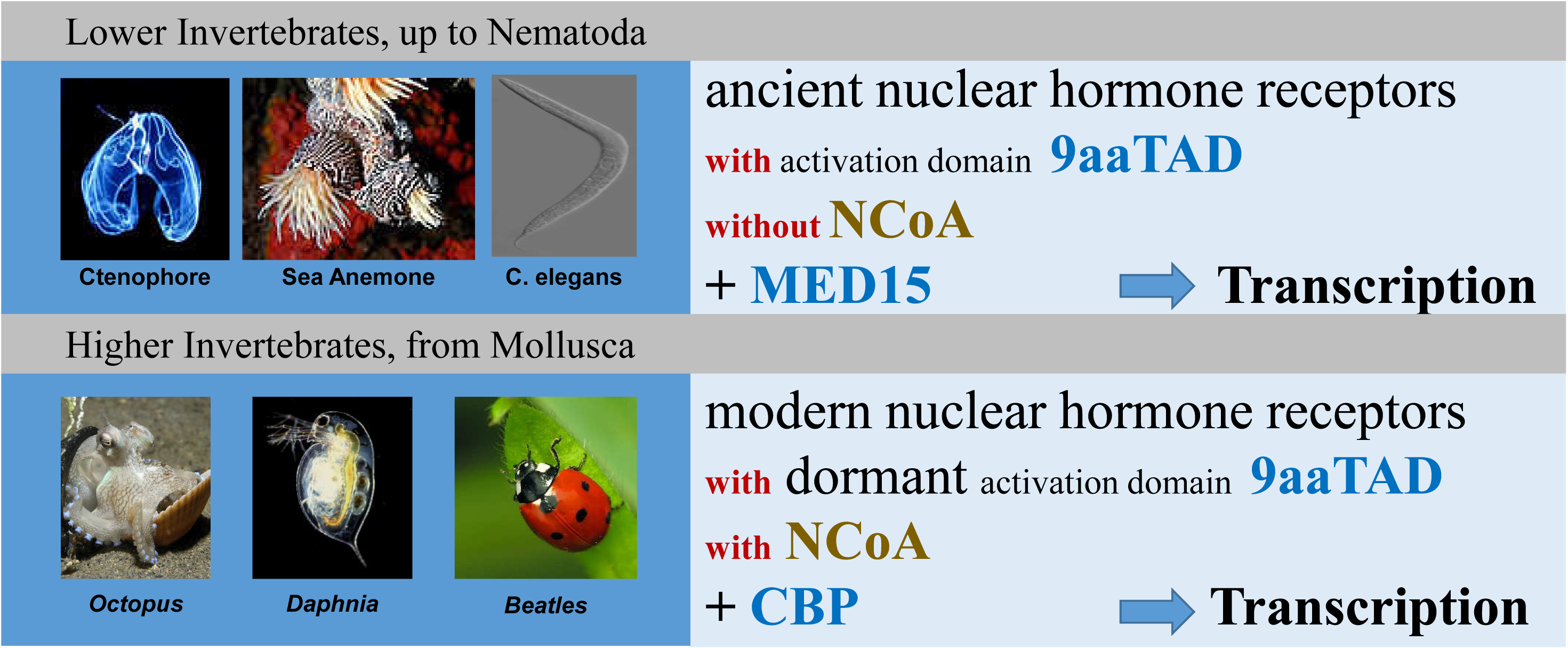

## Introduction

The 9aaTAD activation domains are universal activators of transcription in numerous transcription factors, which are recognized throughout the eukaryotes ^1–3^. The 9aaTAD domains are recognized by multiple mediators of transcription including TAF9, MED15, CBP and p300 ^4–9^. We have also demonstrated that the activation domains 9aaTAD from p53, MLL, E2A, NFkB, Sp1, CTF and SOX transcription factors activated transcription as small peptides ^1,2,10,11^.

The direct interaction of transcription factors and general mediators of transcription is the canonical activation of transcription, which was lost in nuclear hormone receptors in higher eukaryotes and have been replaced by specific hormone adaptors, NCoAs (SRC-1/NCoA-1, TIF-2/NCoA-2 or RAC-3/NCoA-3)^12–14^.

The nematode *C.elegans* has a complex physiology including neuronal network, reproduction organs with ovary in female and testis in male ^15^. Most of nuclear hormone receptors in *C.elegans* have a common HNF4 ancestor and share high degree of homology ^16^. The nuclear hormone receptor NHR-49 is the key regulator of lipid metabolism, dietary response and life span in *C.elegans* 17–19. Both NHR-49 and NHR-64 nuclear hormone receptors are recognized directly by the transcriptional mediator MED15 (MDT-15) ^17^.

Recently from the Taubert lab ^20^, the 9aaTAD domain was identified in the nuclear hormone receptor NHR-49 by our 9aaTAD prediction algorithm (www.piskacek.org). The gain-of-function mutation NHR with V411E substitution was shown to cause significant reduction in MED15 binding ^20^.

In this study, we investigated the 9aaTAD function in *C.elegans* nuclear hormone receptor NHR-49 in parallel to its human homolog Hepatocyte Nuclear Factor 4 (HNF4) and paralogs including Retinoic acid receptor alpha (RARa), Vitamin D receptor (VDR) and Peroxisome proliferator-activated receptor gamma (PPARg).

## Results

### Activation and Inhibitory domains in the nuclear hormone receptor NHR-49

The nuclear hormone receptor NHR-49 in *C.elegans* interacts with transcriptional mediator MED15 (MDT-15) ^17^. The mutation V411E in NHR-49 caused significantly reduced MED15 binding.

We generated hybrid constructs with LexA (prokaryotic DNA binding domain) and activation domain of the NHR-49 (**Figure 1A**). The construct NHR-49 (H11+H12+F, 391-438 aa) included the activation domain 9aaTAD (located in the H12 domain) and adjacent domains H11 and F. The construct (H11+H12, 391-424 aa) included the 9aaTAD activation domain and the adjacent domain H11.

**Figure 1.**
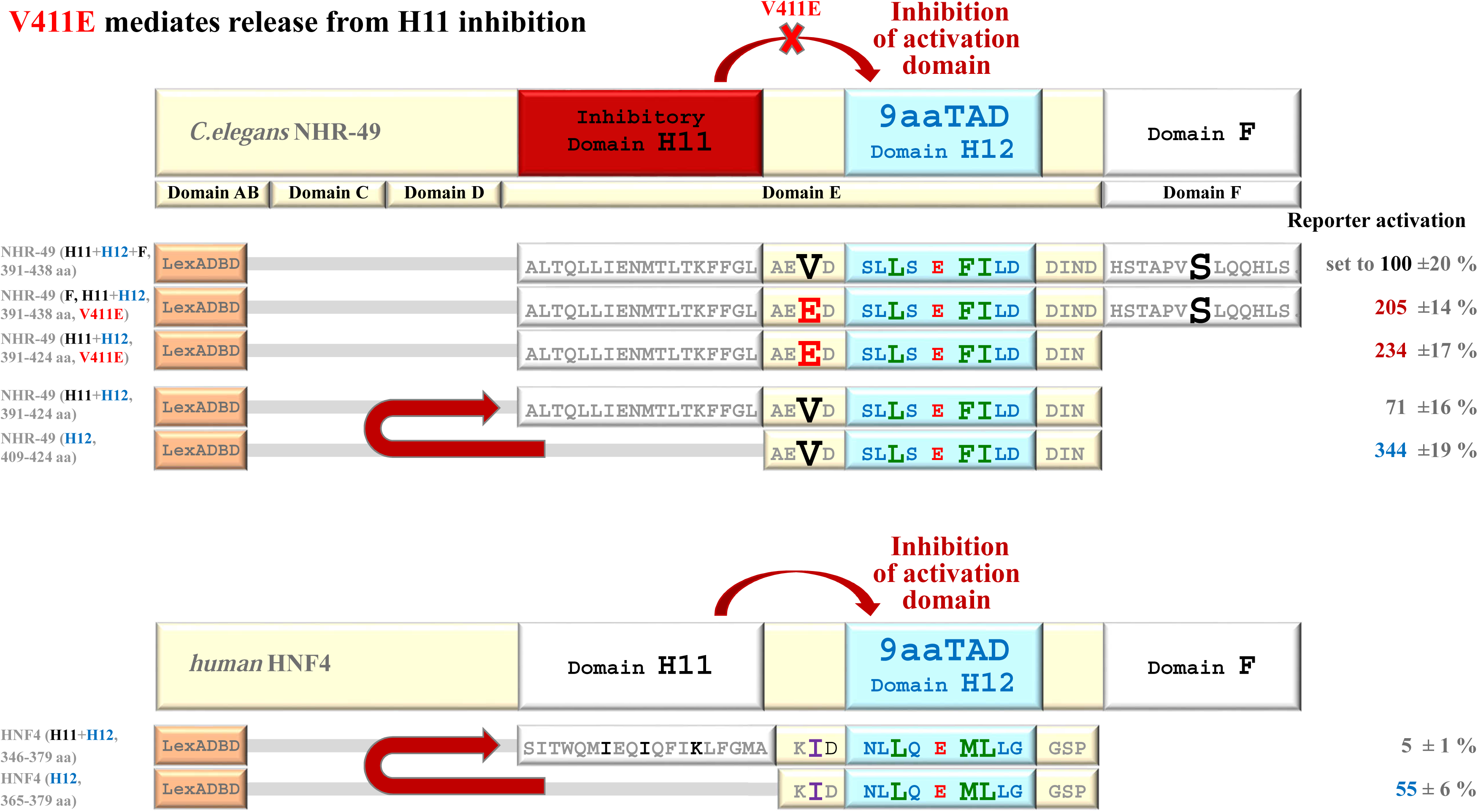
Predicted activation domains 9aaTAD in the C.elegans NHR-49 and human HNF4a nuclear hormone receptors are activators of transcription. The presence inhibitory domains and interference of NHR-49 V411E with the inhibition was monitored. The DNA bindingdomain of LexA were used with parts of NHR-49 for generation of hybrid constructs (BTM116 backbone, standard LexA hybrid assay with β-galactosidase reporter). The average values of the β-galactosidase activities from duplicates experiments are presented as a percentage of the reference with standard deviation (substrate ONPG, means and plusmn; SD; n = 3). We standardized all results to the construct NHR-49 F, including activation domain 9aaTAD and both adjacent domains H11 and F, which activity was setto 100%. The construct DD with deleted Gal4 activation domain served as negative control (sequence shown in Figure 3).

The construct NHR-49 (H12, 409-424 aa) included only the activation domain 9aaTAD. The constructs NHR-49 (H11+H12+F, 391-438 aa, V411E) and NHR-49 (H11+H12, 391-424 aa, V411E) included the mutation V411E.

The reporter activation was monitored by the β-galactosidase assay (LexA dependent reporter for transcription activation assay). We standardized all results in **Figure 1** to the construct NHR-49 (H11+H12+F, 391-438 aa, V411E) and set the activity to 100%. The expression of the proteins were confirmed for each constructs by western blotting (**Figure S1**). The constructs with the mutation V411E showed significantly higher reporter activation than the corresponding wild type construct. Therefore, we concluded that the mutation V411E interfered with inhibitory domain activity.

The construct NHR-49 (H12) including only the 9aaTAD domain showed the most activity in contrast to constructNHR-49 (H11), which included both the activation domain 9aaTAD and the adjacent domain H11. These results suggested the presence of an inhibitory domain within the H11 region.

### Activation and Inhibitory domains in human nuclear hormone receptor NHF4

Following the observations in nuclear hormone receptor NHR-49 in *C.elegans*, we next investigated the human HNF4 receptor activation domain and its adjacent domain H11. Several HNF4 constructs were reported previously ^21^ (**Figure S2**), some that included the 9aaTAD domain and activated transcription and other constructs that have not.

Similarly as above, we predicted the presence of inhibitory domain, which could inhibit the 9aaTAD activation domain in HNF4. Accordingly, we generated LexA constructs with and without the predicted inhibitory H11 domain. The construct HNF4 (H11+H12, 346-379 aa) included the activation domain and the adjacent H11 domain. The construct HNF4 (H12, 365-379aa) included only the activation domain 9aaTAD located in H12 region (**Figure 1B**).

We then compared the construct HNF4 (H12) to a very strong activator Gal4 and found that the activity of the construct HNF4 (H12) reached about 28% of the Gal4 activity (**Figure 2**). The activation domain I and II of the human and rat p53, which all share sequence similarity with hormone receptors, was added to experiment. We concluded that the 9aaTAD activation domain is a powerful activator in HNF4. The HNF4 inhibitory domain inhibited about 10-fold the 9aaTAD function (**Figure 2**). Our observations and conclusions are in a good agreement with previously reported HNF4 constructs ^21^ (**Supplementary Figure S2**).

**Figure 2.**
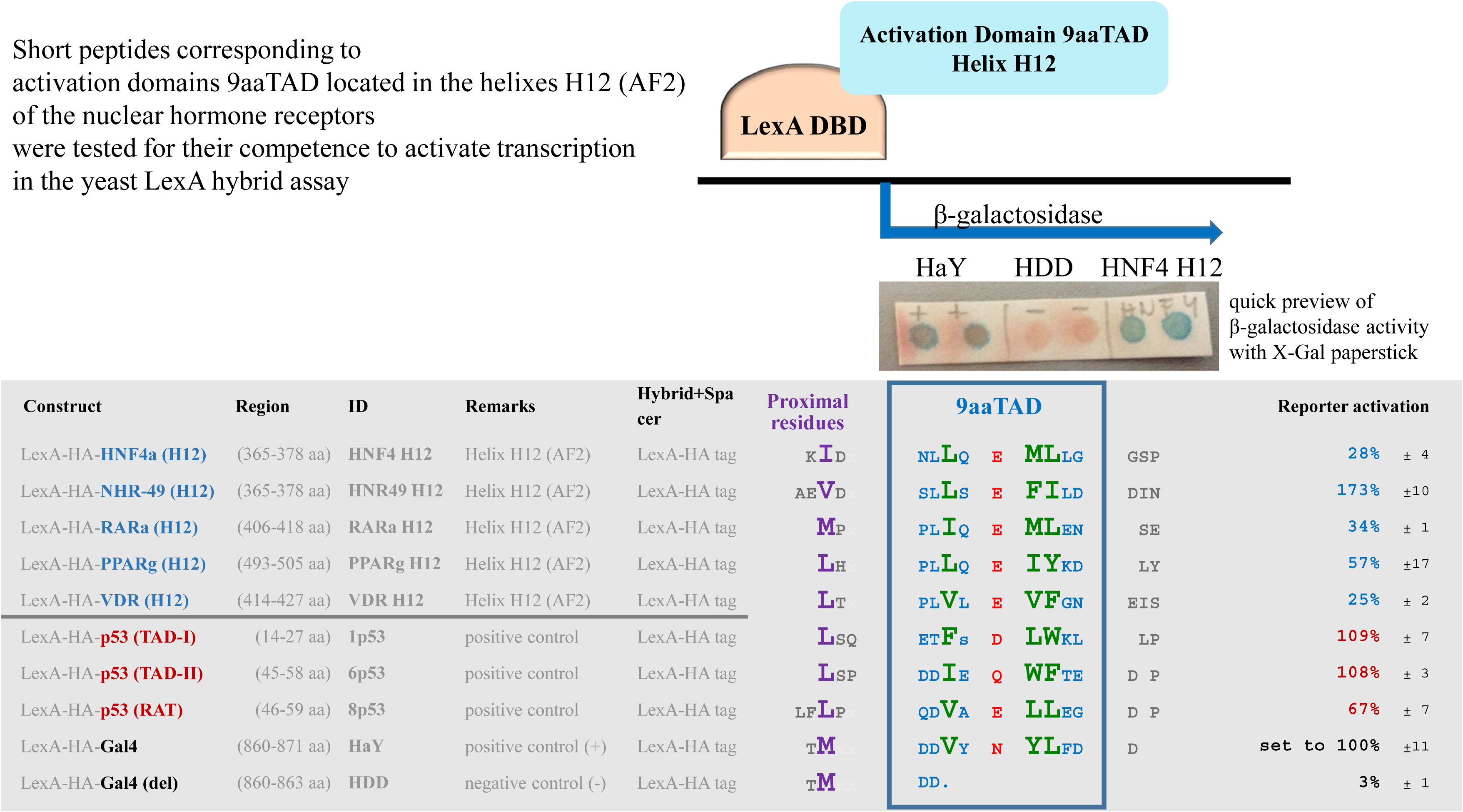
Predicted activation domains 9aaTAD in the human nuclear hormone receptor are universal activators of transcription. The DNA binding domain of LexA were used with putative activation domains for generation of hybrid constructs (BTM116 backbone, standard LexA hybrid assay with β-galactosidase reporter). The average values of the β-galactosidase activities from duplicates experiments are presented as a percentage of the reference with standard deviation (substrate ONPG, means and plusmn; SD; n = 3). The β-galactosidase was previewed by using 0,5 % X-gal on filter paper (no value are available) and show for demonstration. We standardized all results to the previously reported strong Gal4 activator of transcription, construct HaY including LexA and Gal4 activation domain 9aaTAD, which activity was set to 100%. The construct DD with deleted Gal4 activation domain served as negative control. The activity of the LexA-p53 constructs 1p53, 6p53 and 8p53 are shown as extended positive controls and comparable activation domain sequences.

### Activation domains in the human nuclear hormone receptors

Next, we generated sequence alignments (UniProt DB) of selected nuclear hormone receptors including RARa, PPARg and VDR and focused on C-terminal ends with the H10, H11 and H12 domains (described previously for HNF4 and RARa ^22,23^)(**Figure 3**).

**Figure 3.**
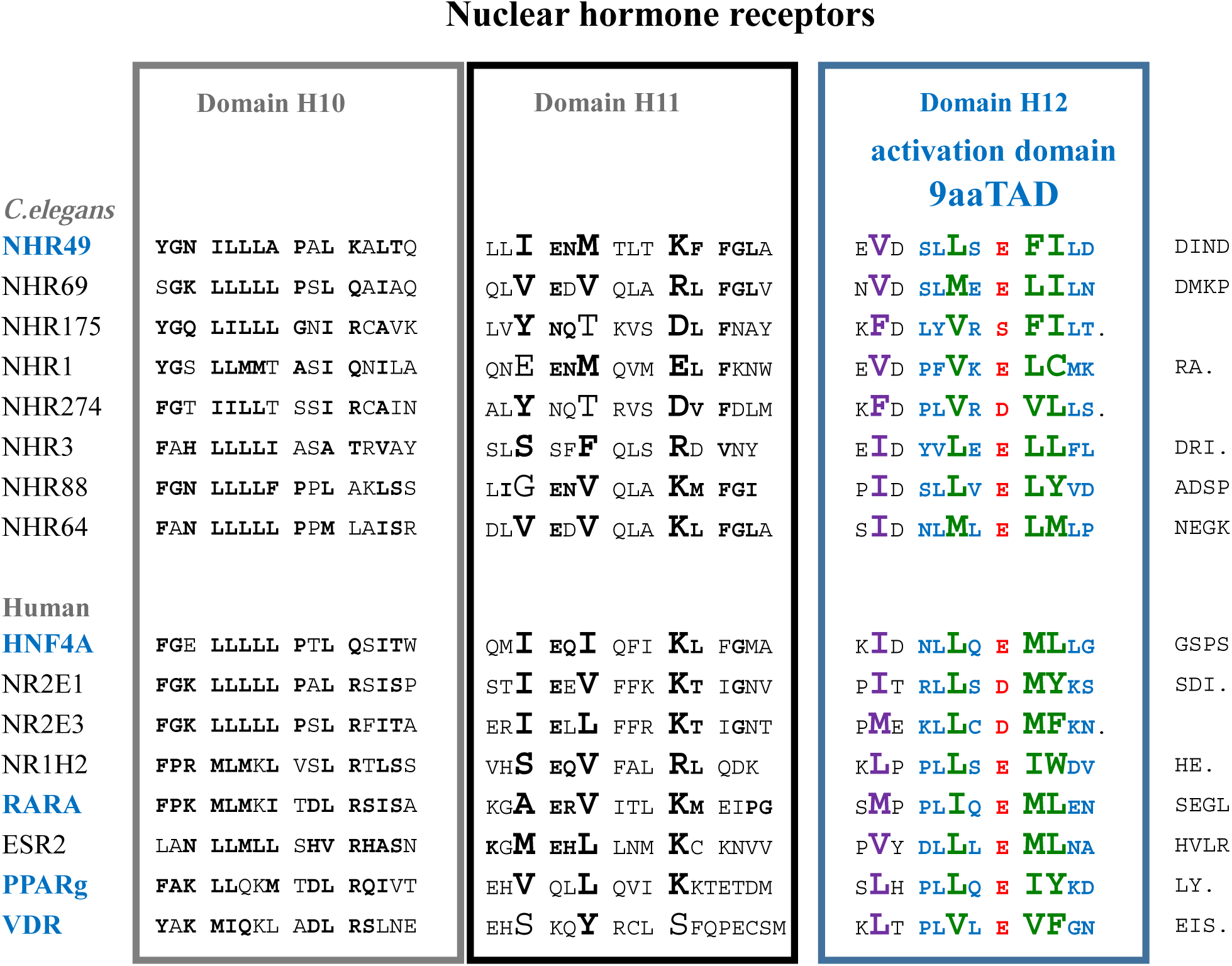
Sequence alignment of the nuclear hormone receptor Family. The C-terminal sequences of selected nuclear hormone receptors from *C.elegans* and humans were aligned. Those activation domains were cloned and tested for activation of transcription(shown in Figure 3) are blue labelled. The structural domains H10, H11 and H12 are clustered. The amino acids in H11, which are like to provide interaction with H12 are bolded.The predicted activation domains 9aaTAD are coloured for fast orientation. The end of sequences are marked with single dots.

We have predicted putative 9aaTAD activation domains in several nuclear hormone receptors including RARa, PPARg and VDR. We then generated LexA constructs with the predicted 9aaTAD activation domains and tested their ability to activate transcription. We then confirmed that all 9aaTADs in RARa, PPARg and VDR hormone receptors are strong activators of transcription as small peptides (**Figure 2**) and that without their hormone adaptors NCoAs.

### Activation and Inhibitory domains in the nuclear hormone receptor RARa

In parallel to NHR-49, we also identified inhibitory domains in RARa, which could cause the inhibition of the predicted activation domain 9aaTAD. Accordingly, we generated LexA constructs with and without the H11 predicted inhibitory domain. The construct RARa (H12) included only the activation domain 9aaTAD located in H12 region. The construct RARa (H11+12) included the H12 region with activation domain 9aaTAD plus the adjacent H11 domain (**Figure 4**). As expected, only the construct RARa (H12) without the inhibitory domain H11 powerfully activated transcription.

**Figure 4.**
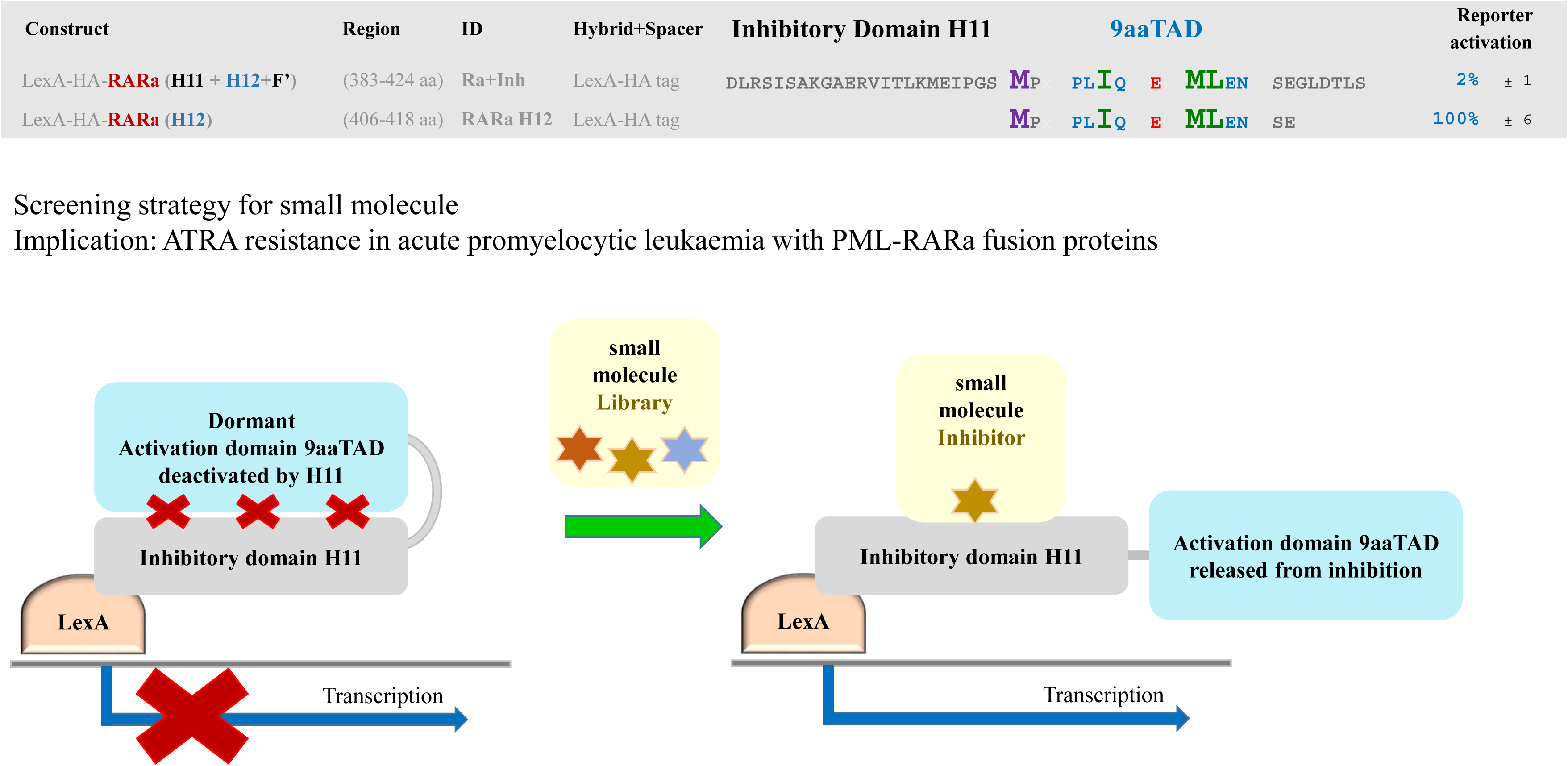
RARa activation and inhibitory domain - Screening strategy for small molecule. The DNA binding domain of LexA were used with RARa activation domains with and without inhibitory domains for generation of hybrid constructs (BTM116 backbone, standard LexA hybrid assay with β-galactosidase reporter). The average values of the β-galactosidase activities from duplicates experiments are presented as a percentage of the reference with standard deviation (substrate ONPG, means and plusmn; SD; n = 3). The Screening strategy for small molecule are schematically shown.

## Discussion

The nuclear hormone receptors in vertebrates are evolutionary distinguished group of unconventional transcription factors using specific adaptors NCoAs, including SRC-1/NCoA-1, TIF-2/NCoA-2 or RAC-3/NCoA-3, to activate transcription ^22,24,25,12–14^.

The NCoA adaptors are essential for nuclear hormone receptor activation in higher invertebrates and all vertebrates, but are completely absent in *C.elegans* genome and all other early branched invertebrata including nematoda, annelida, cnidaria, placozoa, porifera and ctenophora (database Kyoto Encyclopedia of Genes and Genomes, KEGG, Mnemiopsis Genome Project Portal at NHGRI/NIH, genome assembly Ensembl EMBL-EBI)(graphical abstract) ^26^. The absence of the NCoA adaptors in *C.elegans* therefore requires direct interaction of the nuclear hormone receptors with the general mediators of transcription.

In this study, we demonstrated that small 9aaTAD peptides derived from nuclear hormone receptors effectively activate transcription in absence of the NCoA adaptors. We showed that adjacent domains in these hormone receptors have inhibitory effect on their 9aaTAD activation domains.

In the higher eukaryotes, the interface of the nuclear hormone receptor HNF4 with the NCoA adaptors involve multiple regions H3, H4 and H12 domains ^22^. The conserved residua Lys194 (from helix H3) and Glu363 (from helix H12) (**Figure 5 and 6**) together with the hydrophobic cleft created by hydrophobic residua Val190 and Phe199 (from H3 and H4 domains) facilitated NCoA1 adaptor binding.

**Figure 5.**
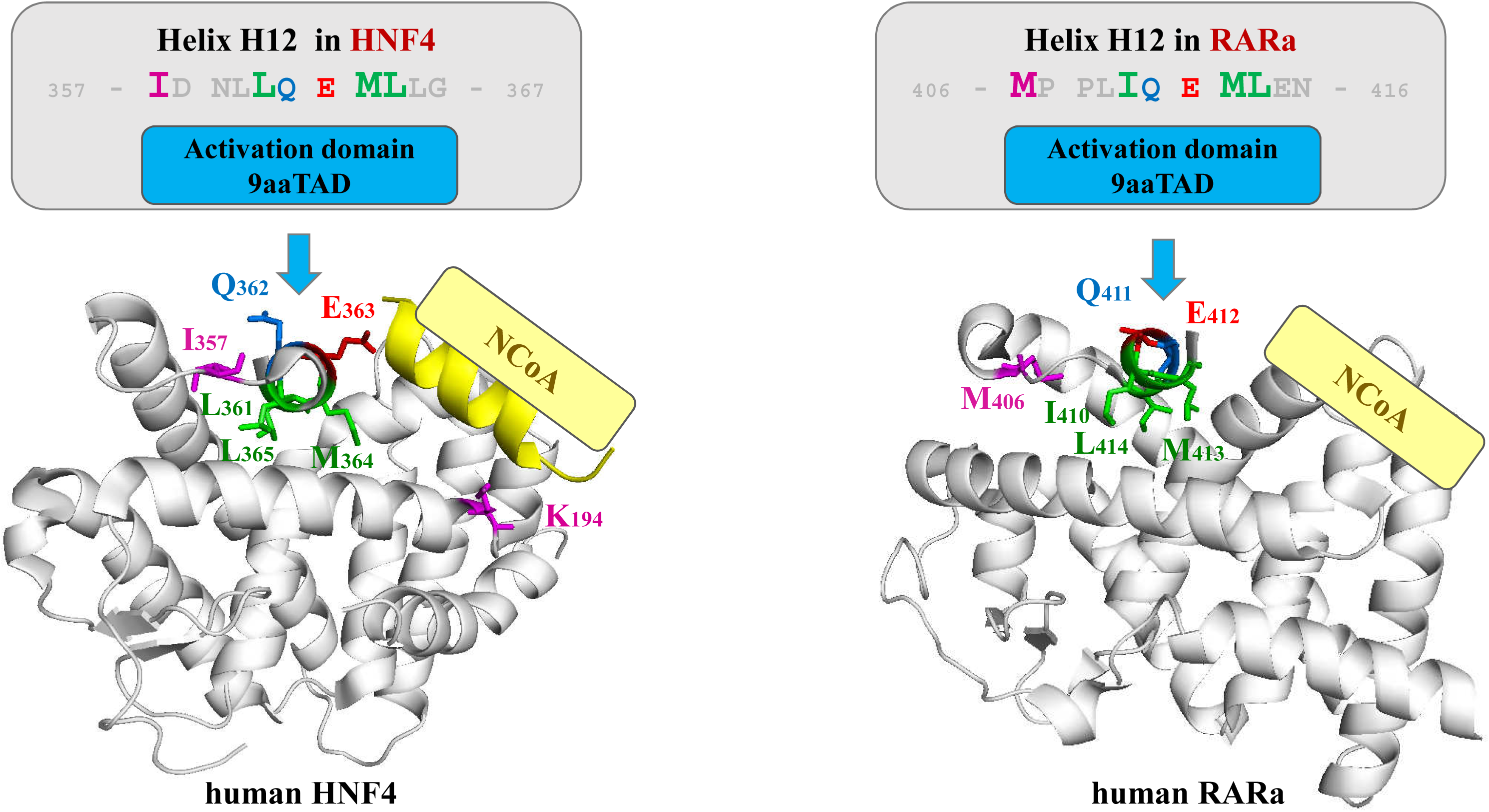
Structures of the human nuclear hormone receptors HNF4a and RARa. The activation domains 9aaTAD are shown and selected residua are coloured. The residue I_357_ in HNF4a, which corresponds to V_411_ in NHR-49, is labelled purple. The HNF4a residue critical for interaction with NCoA E_363_ and K_194_ are labelled red and purple.

**Figure 6.**
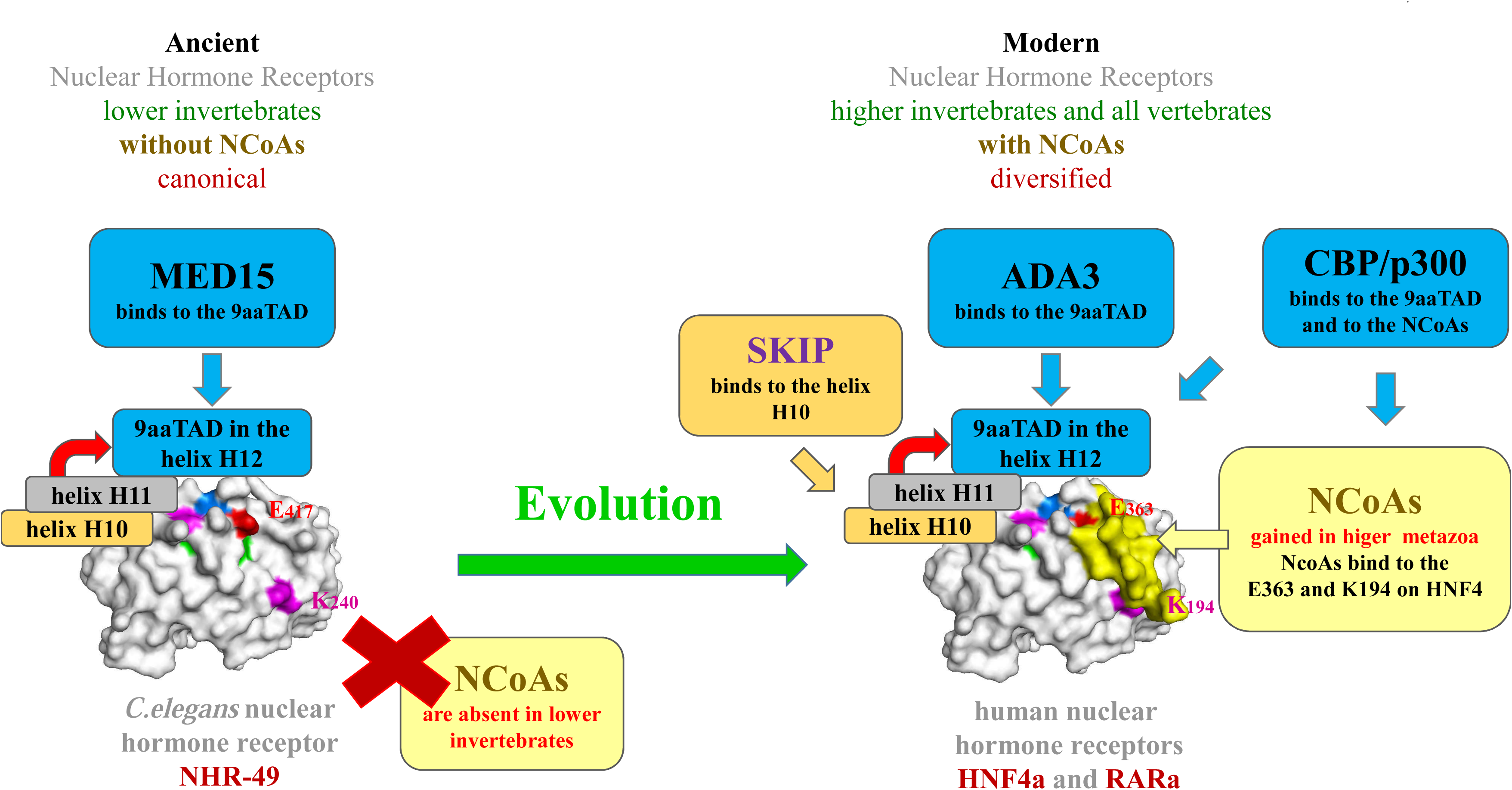
Evolution in the nuclear hormone receptor family. Schema of the conventional(canonical) activation of transcription in ancient nuclear hormone receptor NHR-49, which is strait forward and is provided by general mediator of transcription MED15, but highly complex and diversified in the modern hormone system with specific NCoA adaptors (in yellow). The activation domains 9aaTAD are coloured for fast orientation. For details see text.

In the *C.elegans* NHR-49, where NCoA adaptors are absent, these residua are also conserved (Lys194/240, Glu363/417, Val190/236 and Phe199/245) and might be an evolution prone for the modern nuclear hormone NCoA adaptors, which merged already in mollusca (graphical abstract).

Previously, the bivalent NHF4 character with AF1 and AF2 activation domains were reported ^22^. Strikingly, only a single residue (Glutamic acid Glu363) from the AF2 domain was involved in the interaction with NCoA. The other residua of the HNF4 involved in interaction with NCoA are located far away in the HNF4 protein from the activation domains AF1 and AF2 (Lys194, Val190 and Phe199)(**Figure 5**). The Glu363 residue is in the position p5 of the activation domains 9aaTAD, which generally face to solvent and did not contribute to binding with mediators as are CBP, p300, MED15 ^27^. For the activation domain 9aaTAD function, only the region of the H12 domain is needed (this study). Differently, almost the entire receptor structural integrity is needed for the NCoA mediated activation of transcription.

Following the above, the modern hormone nuclear receptors posses two activation pathways, which one is primordial (canonical, found already in *C.elegans* NHR-49, independent on NCoAs and almost dormant in modern receptors) and another is gained (dependent on NCoAs and required structural integrity of the whole ligand binding domain of the receptor).

The position, structure and motif of the 9aaTAD domain in the nuclear hormone receptor HNF4 matches to human RARa ^22,23^ (**Figure 5**). The H12 helix is in well agreement with other helical activation domains 9aaTAD ^27–29^. The RARa interacts directly with the general mediator ADA3 of transcription also without NCoAs involvement, suggesting an early ancestry ^23,30^. The interaction was restricted by mutants analysis to the H12 region and therefore identical to the 9aaTAD activation domain.

Another interaction, which is outside of the major NCoA scheme for vertebrates nuclear hormone receptors, was reported for human nuclear hormone receptor PPARg. In a ligand-dependent manner, the PPARg interacted directly with mediators p300 and CBP and that without attendance of the NCoA adaptors ^31–34^. The last six amino acids of the PPARg, which are part of the activation domain 9aaTAD, were essential for the interaction. The interaction of CBP or p300 was reported also for other nuclear hormone receptors including PPARa, RARa, RXR, TR and ER ^34–38^. These observations did not oppose the function of molecular adaptors, rather show an alternative for the transcriptional activation in modern nuclear hormone receptors.

The conventional activation of transcription facilitated by direct interaction of activation domains with the general mediators of transcription were found in the most of transcription factors inclusive ancient nuclear hormone receptors. The modern nuclear hormone adaptors NCoA are evolutionary gained and broad out the diversification of the nuclear hormone network. The activation of transcription found in ancient and modern nuclear hormone receptors showed intriguing origin, which differentiated during evolution from simple to complex.

The conservation of the ancestral 9aaTAD activation domains in hormone receptors offered new strategy against cancer. The PML-RARa fusion protein cause about 95% acute promyelocytic leukaemia (APL) and could be well treated in most cases by all-trans retinoic acid (ATRA). However, the ATRA resistance is observed already during initial treatments and is more pronounced during relapses ^39,40^. The mutations of the RARa causing ATRA resistance spread over the ligand binding region in PML-RARa fusions. Therefore, the nuclear hormone adaptors NCoAs could no longer bind to mutated PML-RARa. In the most reported cases, the dormant 9aaTAD activation domain stayed intact (UniProt DB), but is still naturally inactivated by RARa inhibitory domain H11. Thus, a small molecule, which would released the 9aaTAD from the inhibition might provide the cure.

## Materials and Methods

### Constructs

The construct pBTM116-HA was generated by insertion of the HA cassette into the EcoRI site of the vector pBTM116 (HA cassette nucleotide sequence: TGG CTG - GAATTA - GCC ACC ATG GCT TAC CCA TAC GAT GTT CCA GAT TAC GCT GTC GAG ATA - GAATTC, which render in amino acids sequence: W L - E L - A T M A Y P Y D V P D Y A V E I - E F). The constructs were generated by PCR and sub-cloned into the multi-cloning site of pBTM116 vector by EcoRI and PstI sites. All constructs were sequenced by Eurofins Genomics. Further detailed information about constructs, primer sequences are available on the request.

### Assessment of enzyme activities

The β-galactosidase activity was determined in the yeast strain L40 41,42. The strain L40 has integrated the lacZ reporter driven by the lexA operator. In all hybrid assays, we used 2μ vector pBTM116 for generation of the LexA hybrids. The yeast strain L40, the Saccharomyces cerevisiae Genotype: MATa ade2 his3 leu2 trp1 LYS::lexA-HIS3 URA3::lexA-LacZ, is deposited at ATCC (#MYA-3332). For β-galactosidase assays on paper stick, the cells were dropped on Watman paper, lysed by two free/thaw cycle (-20°C), soaked in about 400μl of 100 mM phosphate buffer pH7 with 10 mM KCl, 1 mM MgSO4 and 0.4% X-gal, incubated at 37°C (in a plastic well) until the positive control turned in blue and dried on tissue paper. For standard β-galactosidase assays, overnight cultures propagated in YPD medium (1% yeast extract, 2% bactopeptone, 2% glucose) were diluted to an A600 of 0.3 and further cultivated for two hours and collected by centrifugation. The crude extracts were prepared by vortexing with glass beads for 3 minutes. The assay was done with 10 ul crude extract in 1ml of 100 mM phosphate buffer pH7 with 10 mM KCl, 1 mM MgSO4 and 0.2% 2-Mercaptoethanol; reaction was started by 200 ul 0.4% ONPG and stopped by 500 ul 1 M CaCO3. The average value of the β-galactosidase activities from three independent experiments is presented as a percentage of the reference with the standard deviation (means and plusmn; SD; n = 3). We standardized all results to previously reported Gal4 construct HaY including Gal4 activation domain 9aaTAD with the activity set to 100% 1.

### Western Blot Analysis

The crude cell extracts were prepared in a buffer containing 200 mM Tris-HCl, pH 8.0, 1 mM EDTA, 10% glycerol (v/v), separated by SDS-PAGE, and blotted to nitrocellulose. The immuno-detection of proteins was carried out using mouse anti-HA antibody (#26183, ThermoFisher Sci) or mouse anti-LexA (#306-719, EMD Millipore Corp). The secondary antibodies used were anti-mouse IgG antibodies conjugated with horseradish peroxidase (#A9044, Sigma Aldrich). The proteins were visualized using Pierce ECL (#32106, ThermoFisher Sci) according to the manufacturer’s instructions.

### Source of Photographs

Wikipedia commons download: C. elegans,Zeynep F. Altun, Editor of www.wormatlas.org, https://commons.wikimedia.org/wiki/File:Adult_Caenorhabditis_elegans.jpg, Anemone, Nhobgood, Nick Hobgood, https://upload.wikimedia.org/wikipedia/commons/6/64/Striped_colonial_anemone.jpg, Daphnia pulex,Paul Hebert, doi:10.1371/journal.pbio.0030219, https://commons.wikimedia.org/wiki/File:Daphnia_pulex.png, Octopus,Nick Hobgood, https://commons.wikimedia.org/wiki/File:Octopus_shell.jpg, Drosophila,André Karwath aka Aka, https://commons.wikimedia.org/wiki/File:Drosophila_melanogaster_-_side_(aka).jpg, Seven-spot ladybird,Dominik Stodulski, Graphic Processing: Math Knight, https://commons.wikimedia.org/wiki/File:7-Spotted-Ladybug-Coccinella-septempunctata-sq1.jpg, Ctenophore,Image courtesy of Arctic Exploration 2002, Marsh Youngbluth, NOAA/OER, https://commons.wikimedia.org/wiki/File:Bathocyroe_fosteri.jpg.

## Acknowledgments

This work was supported by the Ministry of Health of the Czech Republic 15-32935A.

---

**Suppl. Figure S1.**
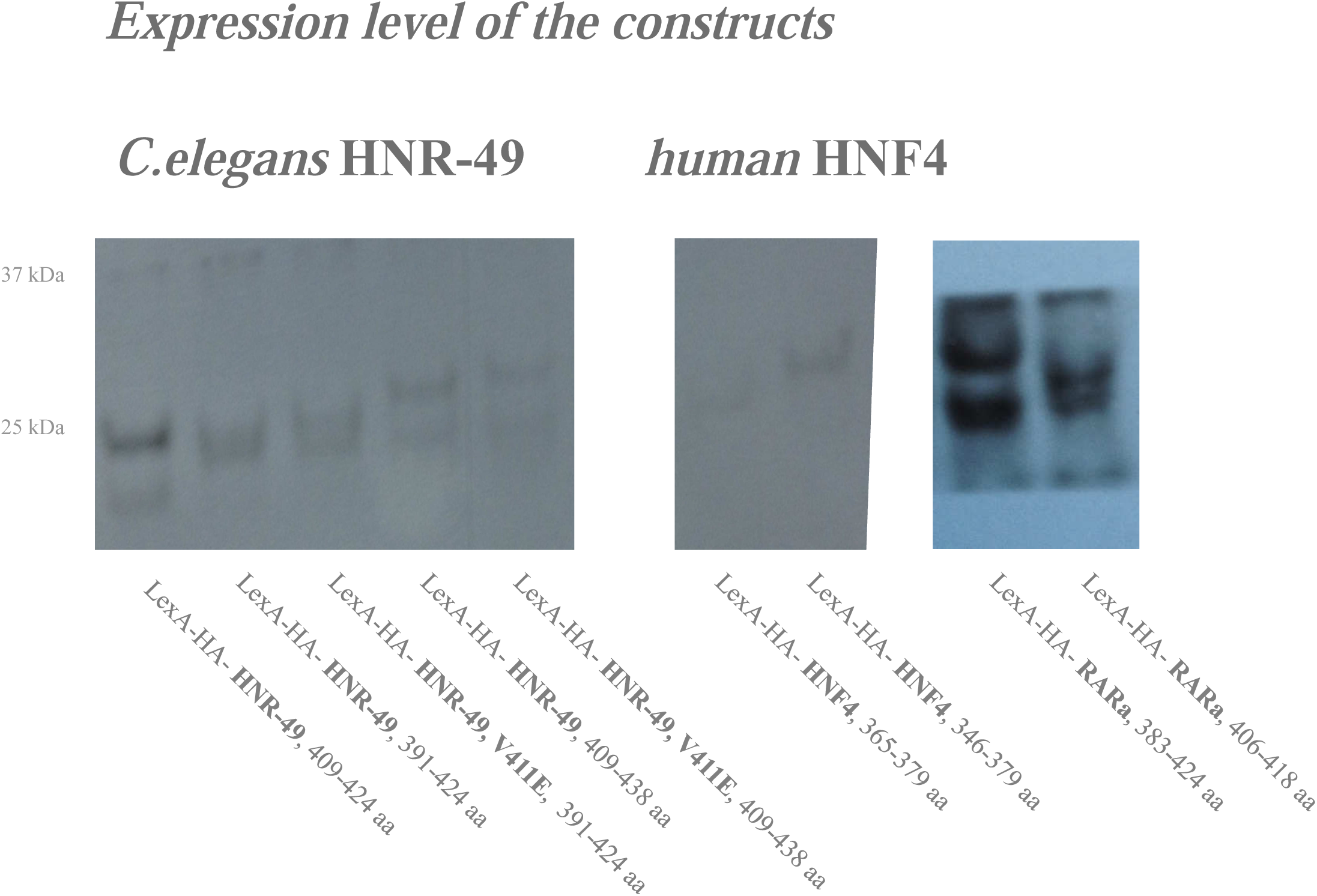
Expression of the constructs. The protein level produced from the constructs were monitored by Western blotting. The proteins comprise of LexA DNA binding domain. The peptides shown were tested in the reporter assay with hybrid Lex DNA binding domain for the capacity to activate transcription.

**Suppl. Figure S2.**
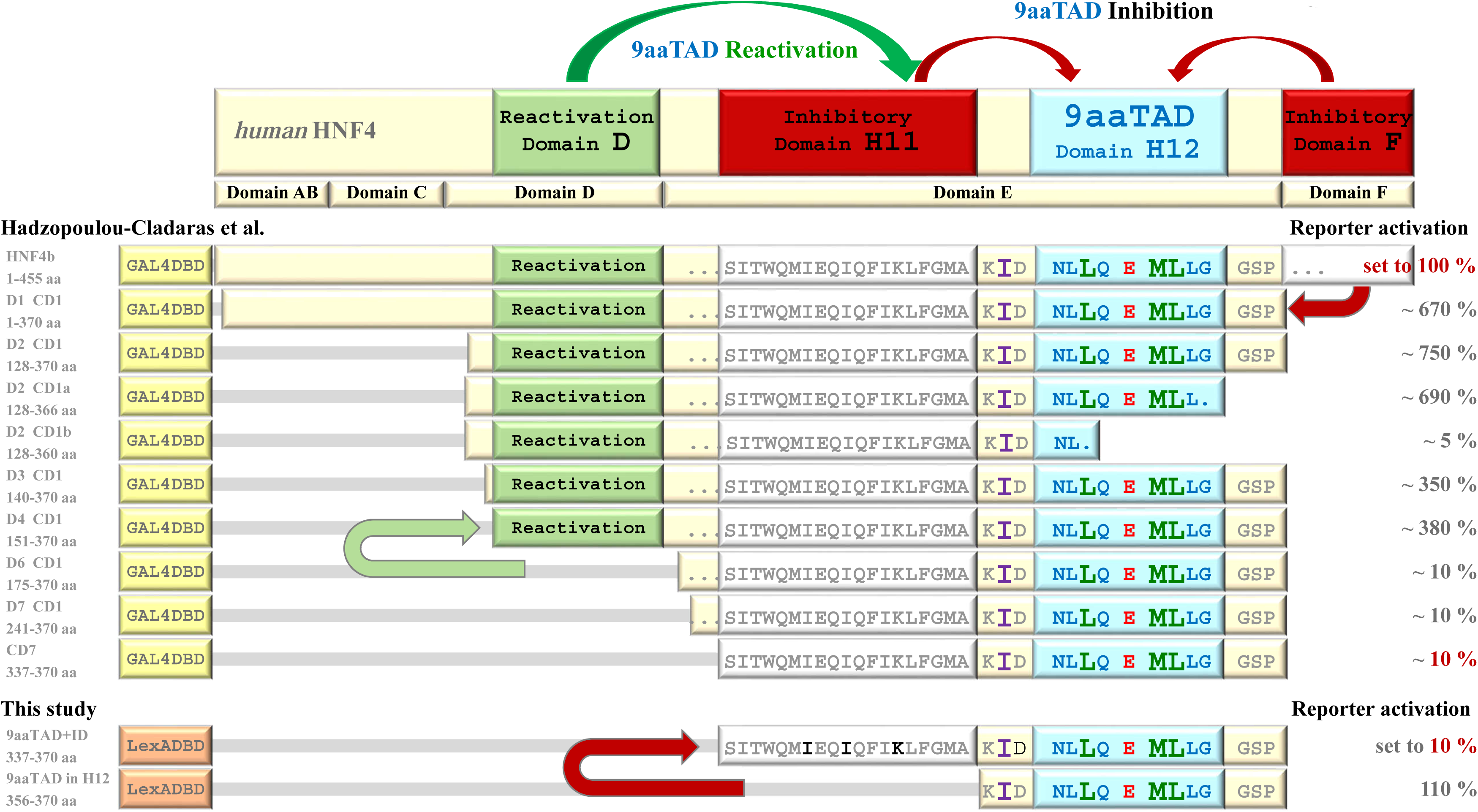
Activation of transcription by HNF4 constructs. The schema of human HNF4 constructs from Hadzopoulou-Cladaras et al. 1997 and our constructs from Figure 1 are aligned and their functional domains are graphically organised according structural domain H11 and H12, originally domain A to F and activation of transcription are shown in percent. Our HNF4 construct 9aaTAD+ID correspond to reported construct CD7 and therefore correspond to activity of 10%. The activation domains 9aaTAD are colouredfor fast orientation. The proximal amino acid I357 corresponding to V411 in C.elegans NHR-49 is in purple. The arrows indicated two lost of functional domain betwenn two constructs.

## References

1 Piskacek M, Havelka M, Rezacova M, Knight A. The 9aaTAD Transactivation Domains: From Gal4 to p53. PLoS ONE 2016; 11: e0162842.

2 Piskacek M, Havelka M, Rezacova M, Knight A. The 9aaTAD Is Exclusive Activation Domain in Gal4. PLoS ONE 2017; 12: e0169261.

3 Kakidani H, Ptashne M. GAL4 activates gene expression in mammalian cells. Cell 1988; 52: 161–167.

4 Lee CW, Martinez-Yamout MA, Dyson HJ, Wright PE. Structure of the p53 transactivation domain in complex with the nuclear receptor coactivator binding domain of CREB binding protein. Biochemistry 2010; 49: 9964–9971.

5 Thakur JK, Arthanari H, Yang F, Chau KH, Wagner G, Näär AM. Mediator subunit Gal11p/MED15 is required for fatty acid-dependent gene activation by yeast transcription factor Oaf1p. J Biol Chem 2009; 284: 4422–4428.

6 Teufel DP, Freund SM, Bycroft M, Fersht AR. Four domains of p300 each bind tightly to a sequence spanning both transactivation subdomains of p53. Proc Natl Acad Sci USA 2007; 104: 7009–7014.

7 Ma J, Ptashne M. A new class of yeast transcriptional activators. Cell 1987; 51: 113–119.

8 Uesugi M, Verdine GL. The alpha-helical FXXPhiPhi motif in p53: TAF interaction and discrimination by MDM2. Proc Natl Acad Sci USA 1999; 96: 14801–14806.

9 Di Lello P, Jenkins LMM, Jones TN, Nguyen BD, Hara T, Yamaguchi H et al. Structure of the Tfb1/p53 complex: Insights into the interaction between the p62/Tfb1 subunit of TFIIH and the activation domain of p53. Mol Cell 2006; 22: 731–740.

10 Piskacek S, Gregor M, Nemethova M, Grabner M, Kovarik P, Piskacek M. Nine-amino-acid transactivation domain: establishment and prediction utilities. Genomics 2007; 89: 756–768.

11 Sandholzer J, Hoeth M, Piskacek M, Mayer H, de Martin R. A novel 9-amino-acid transactivation domain in the C-terminal part of Sox18. Biochem Biophys Res Commun 2007; 360: 370–374.

12 Heery DM, Kalkhoven E, Hoare S, Parker MG. A signature motif in transcriptional co-activators mediates binding to nuclear receptors. Nature 1997; 387: 733–736.

13 Oñate SA, Tsai SY, Tsai MJ, O’Malley BW. Sequence and characterization of a coactivator for the steroid hormone receptor superfamily. Science 1995; 270: 1354–1357.

14 Voegel JJ, Heine MJ, Zechel C, Chambon P, Gronemeyer H. TIF2, a 160 kDa transcriptional mediator for the ligand-dependent activation function AF-2 of nuclear receptors. EMBO J 1996; 15: 3667–3675.

15 Grants JM, Goh GYS, Taubert S. The Mediator complex of Caenorhabditis elegans: insights into the developmental and physiological roles of a conserved transcriptional coregulator. Nucleic Acids Res 2015; 43: 2442–2453.

16 Taubert S, Ward JD, Yamamoto KR. Nuclear hormone receptors in nematodes: evolution and function. Mol Cell Endocrinol 2011; 334: 49–55.

17 Taubert S, Van Gilst MR, Hansen M, Yamamoto KR. A Mediator subunit, MDT-15, integrates regulation of fatty acid metabolism by NHR-49-dependent and -independent pathways in C. elegans. Genes Dev 2006; 20: 1137–1149.

18 Hibshman JD, Hung A, Baugh LR. Maternal Diet and Insulin-Like Signaling Control Intergenerational Plasticity of Progeny Size and Starvation Resistance. PLoS Genet 2016; 12: e1006396.

19 Van Gilst MR, Hadjivassiliou H, Jolly A, Yamamoto KR. Nuclear hormone receptor NHR-49 controls fat consumption and fatty acid composition in C. elegans. PLoS Biol 2005; 3: e53.

20 Lee K, Goh GYS, Wong MA, Klassen TL, Taubert S. Gain-of-Function Alleles in Caenorhabditis elegans Nuclear Hormone Receptor nhr-49 Are Functionally Distinct. PLoS ONE 2016; 11: e0162708.

21 Hadzopoulou-Cladaras M, Kistanova E, Evagelopoulou C, Zeng S, Cladaras C, Ladias JA. Functional domains of the nuclear receptor hepatocyte nuclear factor 4. J Biol Chem 1997; 272: 539–550.

22 Duda K, Chi Y-I, Shoelson SE. Structural basis for HNF-4alpha activation by ligand and coactivator binding. J Biol Chem 2004; 279: 23311–23316.

23 Li C-W, Ai N, Dinh GK, Welsh WJ, Chen JD. Human ADA3 regulates RARalpha transcriptional activity through direct contact between LxxLL motifs and the receptor coactivator pocket. Nucleic Acids Res 2010; 38: 5291–5303.

24 le Maire A, Bourguet W. Retinoic acid receptors: structural basis for coregulator interaction and exchange. Subcell Biochem 2014; 70: 37–54.

25 Sladek FM. What are nuclear receptor ligands? Mol Cell Endocrinol 2011; 334: 3–13.

26 Hultqvist G, Åberg E, Camilloni C, Sundell GN, Andersson E, Dogan J et al. Emergence and evolution of an interaction between intrinsically disordered proteins. Elife 2017; 6. doi:10.7554/eLife.16059.

27 Piskacek M, Vasku A, Hajek R, Knight A. Shared structural features of the 9aaTAD family in complex with CBP. Mol Biosyst 2015; 11: 844–851.

28 Ernst P, Wang J, Huang M, Goodman RH, Korsmeyer SJ. MLL and CREB bind cooperatively to the nuclear coactivator CREB-binding protein. Mol Cell Biol 2001; 21: 2249–2258.

29 Denis CM, Chitayat S, Plevin MJ, Wang F, Thompson P, Li S et al. Structural basis of CBP/p300 recruitment in leukemia induction by E2A-PBX1. Blood 2012. doi:10.1182/blood-2012-02-411397.

30 Meng G, Zhao Y, Nag A, Zeng M, Dimri G, Gao Q et al. Human ADA3 binds to estrogen receptor (ER) and functions as a coactivator for ER-mediated transactivation. J Biol Chem 2004; 279: 54230–54240.

31 Gelman L, Zhou G, Fajas L, Raspé E, Fruchart JC, Auwerx J. p300 interacts with the N- and C-terminal part of PPARgamma2 in a ligand-independent and -dependent manner, respectively. J Biol Chem 1999; 274: 7681–7688.

32 Lehmann M, Pirinen E, Mirsaidi A, Kunze FA, Richards PJ, Auwerx J et al. ARTD1-induced poly-ADP-ribose formation enhances PPARγ ligand binding and co-factor exchange. Nucleic Acids Res 2015; 43: 129–142.

33 Mizukami J, Taniguchi T. The antidiabetic agent thiazolidinedione stimulates the interaction between PPAR gamma and CBP. Biochem Biophys Res Commun 1997; 240: 61–64.

34 Dowell P, Ishmael JE, Avram D, Peterson VJ, Nevrivy DJ, Leid M. p300 functions as a coactivator for the peroxisome proliferator-activated receptor alpha. J Biol Chem 1997; 272: 33435–33443.

35 Chakravarti D, LaMorte VJ, Nelson MC, Nakajima T, Schulman IG, Juguilon H et al. Role of CBP/P300 in nuclear receptor signalling. Nature 1996; 383: 99–103.

36 Kamei Y, Xu L, Heinzel T, Torchia J, Kurokawa R, Gloss B et al. A CBP integrator complex mediates transcriptional activation and AP-1 inhibition by nuclear receptors. Cell 1996; 85: 403–414.

37 Smith CL, Oñate SA, Tsai MJ, O’Malley BW. CREB binding protein acts synergistically with steroid receptor coactivator-1 to enhance steroid receptor-dependent transcription. Proc Natl Acad Sci USA 1996; 93: 8884–8888.

38 Bojcsuk D, Nagy G, Balint BL. Inducible super-enhancers are organized based on canonical signal-specific transcription factor binding elements. Nucleic Acids Res 2017; 45: 3693–3706.

39 Zhu H-H, Qin Y-Z, Huang X-J. Resistance to arsenic therapy in acute promyelocytic leukemia. N Engl J Med 2014; 370: 1864–1866.

40 Zhou D-C, Kim SH, Ding W, Schultz C, Warrell RP, Gallagher RE. Frequent mutations in the ligand-binding domain of PML-RARalpha after multiple relapses of acute promyelocytic leukemia: analysis for functional relationship to response to all-trans retinoic acid and histone deacetylase inhibitors in vitro and in vivo. Blood 2002; 99: 1356–1363.

41 Miller JH. Experiments in molecular genetics. Cold Spring Harbor Laboratory, 1972.

42 Baumgartner U, Hamilton B, Piskacek M, Ruis H, Rottensteiner H. Functional analysis of the Zn(2)Cys(6) transcription factors Oaf1p and Pip2p. Different roles in fatty acid induction of beta-oxidation in Saccharomyces cerevisiae. J Biol Chem 1999; 274: 22208–22216.

